# A targeted CRISPR screen identifies ETS1 as a regulator of HIV latency

**DOI:** 10.1101/2024.08.03.606477

**Authors:** Manickam Ashokkumar, Terry L Hafer, Abby Felton, Nancie M. Archin, David M Margolis, Michael Emerman, Edward P Browne

**Affiliations:** Department of Medicine, University of North Carolina at Chapel Hill, Chapel Hill, NC 27599, USA; UNC HIV Cure Center, University of North Carolina at Chapel Hill, Chapel Hill, NC 27599, USA; Division of Basic Sciences, Fred Hutchinson Cancer Center, Seattle, Washington, United States of America; Division of Human Biology, Fred Hutchinson Cancer Center, Seattle, Washington, United States of America; Department of Microbiology and Immunology, University of North Carolina at Chapel Hill, Chapel Hill, NC 27599, USA

**Keywords:** HIV latency, CRISPR/Cas9 genome editing, Transcription Factors, ETS1, Person with HIV (PWH)

## Abstract

Human Immunodeficiency virus (HIV) infection is regulated by a wide array of host cell factors that combine to influence viral transcription and latency. To understand the complex relationship between the host cell and HIV latency, we performed a lentiviral CRISPR screen that targeted a set of host cell genes whose expression or activity correlates with HIV expression. We further investigated one of the identified factors - the transcription factor ETS1 and found that it is required for maintenance of HIV latency in a primary CD4 T cell model. Interestingly, ETS1 played divergent roles in actively infected and latently infected CD4 T cells, with knockout of ETS1 leading to reduced HIV expression in actively infected cells, but increased HIV expression in latently infected cells, indicating that ETS1 can play both a positive and negative role in HIV expression. CRISPR/Cas9 knockout of ETS1 in CD4 T cells from ART-suppressed people with HIV (PWH) confirmed that ETS1 maintains transcriptional repression of the clinical HIV reservoir. Transcriptomic profiling of ETS1-depleted cells from PWH identified a set of host cell pathways involved in viral transcription that are controlled by ETS1 in resting CD4 T cells. In particular, we observed that ETS1 knockout increased expression of the long non-coding RNA MALAT1 that has been previously identified as a positive regulator of HIV expression. Furthermore, the impact of ETS1 depletion on HIV expression in latently infected cells was partially dependent on MALAT1. Overall, these data demonstrate that ETS1 is an important regulator of HIV latency and influences expression of several cellular genes, including MALAT1, that could have a direct or indirect impact on HIV expression.

**Author Summary:** HIV latency is a major obstacle for the eradication of HIV. However, molecular mechanisms that restrict proviral expression during therapy are not well understood. Identification of host cell factors that silence HIV would create opportunities for targeting these factors to reverse latency and eliminate infected cells. Our study aimed to explore mechanisms of latency in infected cells by employing a lentiviral CRISPR screen and CRISPR/Cas9 knockout in primary CD4 T cells. These experiments revealed that ETS1 is essential for maintaining HIV latency in primary CD4 T cells and we further confirmed ETS1’s role in maintaining HIV latency through CRISPR/Cas9 knockout in CD4 T cells from antiretroviral therapy (ART)-suppressed individuals with HIV. Transcriptomic profiling of ETS1-depleted cells from these individuals identified several host cell pathways involved in viral transcription regulated by ETS1, including the long non-coding RNA MALAT1. Overall, our study demonstrates that ETS1 is a critical regulator of HIV latency, affecting the expression of several cellular genes that directly or indirectly influence HIV expression.

## Introduction

Despite the development of successful antiretroviral therapy (ART) that can reduce circulating human immunodeficiency virus (HIV) plasma viral loads to undetectable levels, the virus rebounds rapidly from a highly stable latent proviral reservoir following ART interruption (1). Latently infected cells with HIV are thus a major obstacle for the eradication of virus from people with HIV (PWH). Toward this goal, several different classes of latency reversing agents (LRAs) have been developed and investigated for their ability to reactivate latently infected cells and promote their killing by the immune system or by viral cytopathic effect (2-4). However, none of the current LRAs that have been tested *in vivo* have proven to be effective in reducing the size of the HIV reservoir. The reasons for this lack of an impact on the clinical reservoir are unclear, but it is likely the result of the complex relationship between the virus and the set of cellular transcription factors (TFs) that regulate HIV. Individual LRAs that target single pathways are typically inefficient, reflecting the heterogeneous nature of the reservoir, and suggesting that combinations of LRAs will likely be required to achieve broad viral reactivation.

The HIV-1 long terminal repeat (LTR) contains DNA binding sites for several ubiquitously expressed and regulated cellular TFs with activating or repressive functions (5). In recent years, transcriptional regulators (activators or repressors) have been identified which interact directly or indirectly with the HIV-1 5’LTR, and the combined activity of these factors determines the fate of HIV transcription and latency. Nevertheless, the regulation of the viral promoter is still incompletely understood and additional regulatory factors likely exist. Identification of these factors and the characterization of their roles in the maintenance of the HIV reservoir will likely lead to the development of new tools for clinical latency reversal.

We have recently used combined single cell multi-omic profiling of a primary CD4 T cell model of HIV latency to identify a set of host cell genes whose expression or activity correlates with HIV RNA levels (6). Based on this observation we hypothesized that some of these HIV correlated factors could represent novel regulators of HIV expression. Genome editing platforms based on the clustered regularly interspaced palindromic repeat (CRISPR)/CRISPR associated protein 9 (Cas9) system have been recognized as promising tools for probing interactions between the HIV-1 proviral genome and the host cell. Several studies using CRISPR/Cas9 technology have previously identified cellular factors that affect HIV replication or latency (7-10). In this study we have used this set of HIV correlated genes to conduct a targeted CRISPR screen for novel latency regulating factors. Specifically, we have carried out a lentiviral CRISPR-Cas9 pooled knockout screen in J-Lat 10.6 and 5A8 cells, Jurkat cell-based models of HIV-1 latency. This approach revealed several novel genes involved in latency, including the transcriptional regulator, ETS1 (E26 transformation–specific-1). In further validation experiments in primary CD4 T cells and in cells from people with HIV-1, we found that ETS1 has a complex role in establishing latency and reactivation, playing a positive role in actively infected CD4 T cells, but a repressive role in latently infected CD4 T cells.

## Results

### Targeted CRISPR screen design

We have previously shown that HIV expression in CD4 T cells is correlated with expression of a set of host cell genes (6). We hypothesized that a subset of these genes could also be important for controlling HIV expression and latency. Thus, we sought to examine the role of these HIV-correlated genes (**Table S1**) using a previously established HIV-CRISPR screening system (11-13). A schematic description of the screening approach is shown in **Figure 1**. This screening approach uses pooled lentiviral sgRNAs against a set of target genes in two Jurkat-derived cell lines with latent proviruses that encode the HIV Gag protein (J-Lat10.6 and J-Lat5A8). A key aspect of the HIV-CRISPR screen is that the lentiviral guide RNAs (gRNAs) contain an HIV psi (ψ) packaging signal and are thus packaged *in trans* into virions in cells with reactivated HIV gene expression. Thus, if a gene knockout affects viral gene expression, this sgRNA will be either depleted or enriched in the supernatant of the J-Lat cells. We first generated an sgRNA library targeting these 351 genes of interest, as well as a set of positive and negative controls, hereafter referred to as the TxLatent library. The target set for this screen was highly enriched for cellular transcriptional regulators. These guide RNAs (8 per gene along with 150 non-targeting guides) were cloned into an HIV-CRISPR vector, transfected as a pool into HEK293T cells to generate stocks of the HIV-CRISPR particles, then used to transduce two different J-Lat cell lines. After puromycin selection for transduction by the HIV-CRISPR vector, these cells were stimulated in parallel with limiting concentrations of four different latency reversing agents (LRAs) or with an equivalent volume of DMSO to look for synergistic or dependent relationships between these compounds and individual knockouts. The LRAs used were AZD5582 (non-canonical NF-κB agonist), prostratin (protein kinase C agonist), vorinostat (histone deacetylase inhibitor) and iBET151 (bromo-and extra-terminal domain inhibitor) (6). After 24h of LRA stimulation, we harvested genomic DNA and viral supernatants to assess how each gene knockout affected viral production by next-generation sequencing. We validated that there is reactivation of J-Lat lines upon LRA treatment by performing a reverse transcriptase (RT) assay of the two J-Lat lines (Figure S1).

**Figure 1.**
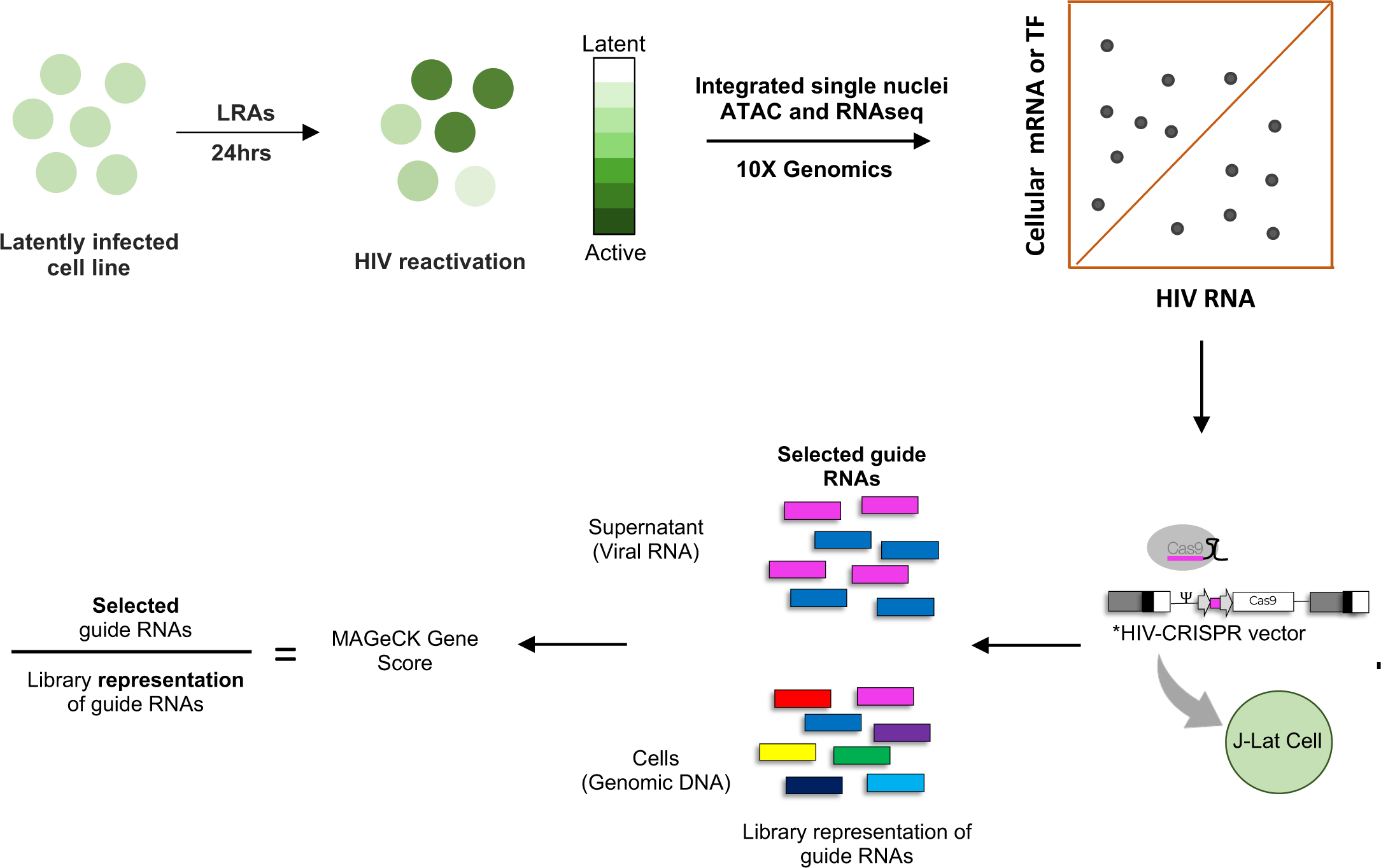
Schematic of targeted CRISPR screen for novel latency regulating factors. In this study we conduct a targeted CRISPR screen for novel latency regulating factors. HIV correlated factors were first identified using a prior dataset and a 351 target pooled sgRNA library (TxLatent) was constructed using the HIV-CRISPR vector that contains an HIV RNA packaging signal. Screening was carried out in J-Lat 10.6 and J-Lat 5A8 cells, two Jurkat cell-based models of HIV-1 latency. sgRNAs that reactivate or inhibit HIV expression are enriched or depleted in secreted virions found in the cellular supernatant respectively. Screening was also carried out in the presence or absence of four different latency reversing agents (LRAs).

We assessed the degree that guides aggregated by gene are either enriched or depleted relative to the non-targeting controls (NTCs) by determining the Model-based Analysis of Genome-wide CRISPR-Cas9 Knockout (MAGeCK) scores of each gene (14) (**Figure 2A-E, Table S2**). We primarily focused on J-Lat 10.6 cells, given these cells showed greater separation from NTCs than in the J-Lat 5A8 cell line (**Table S2, Figure S2**). First, we examined the depleted genes whose knockout prevented viral gene expression during latency reversal in J-Lat 10.6 cells. Consistent with a previous targeted CRISPR screen using this system, *CCNT1*, which encodes Cyclin T1, (**Figure 2B-E**) was among the most depleted targets in J-Lat cells treated with an LRA (13). *CCNT1* knockout inhibited latency reversal under all conditions tested. Similarly, *IKBKB* (the inhibitor of NF-κB Kinase) is a positive regulator of the NF-κB signaling pathway and is also consistently depleted in each LRA treatment (**Figure 2C-E**). Curiously, guides targeting *HDAC3* were also depleted in some conditions (AZD5582; **Figure 2B** and vorinostat; **Figure 2E**), but not in others. Overall, these data confirm that the library and screen design could reliably identify HIV-1 regulating factors in these cells.

**Figure 2.**
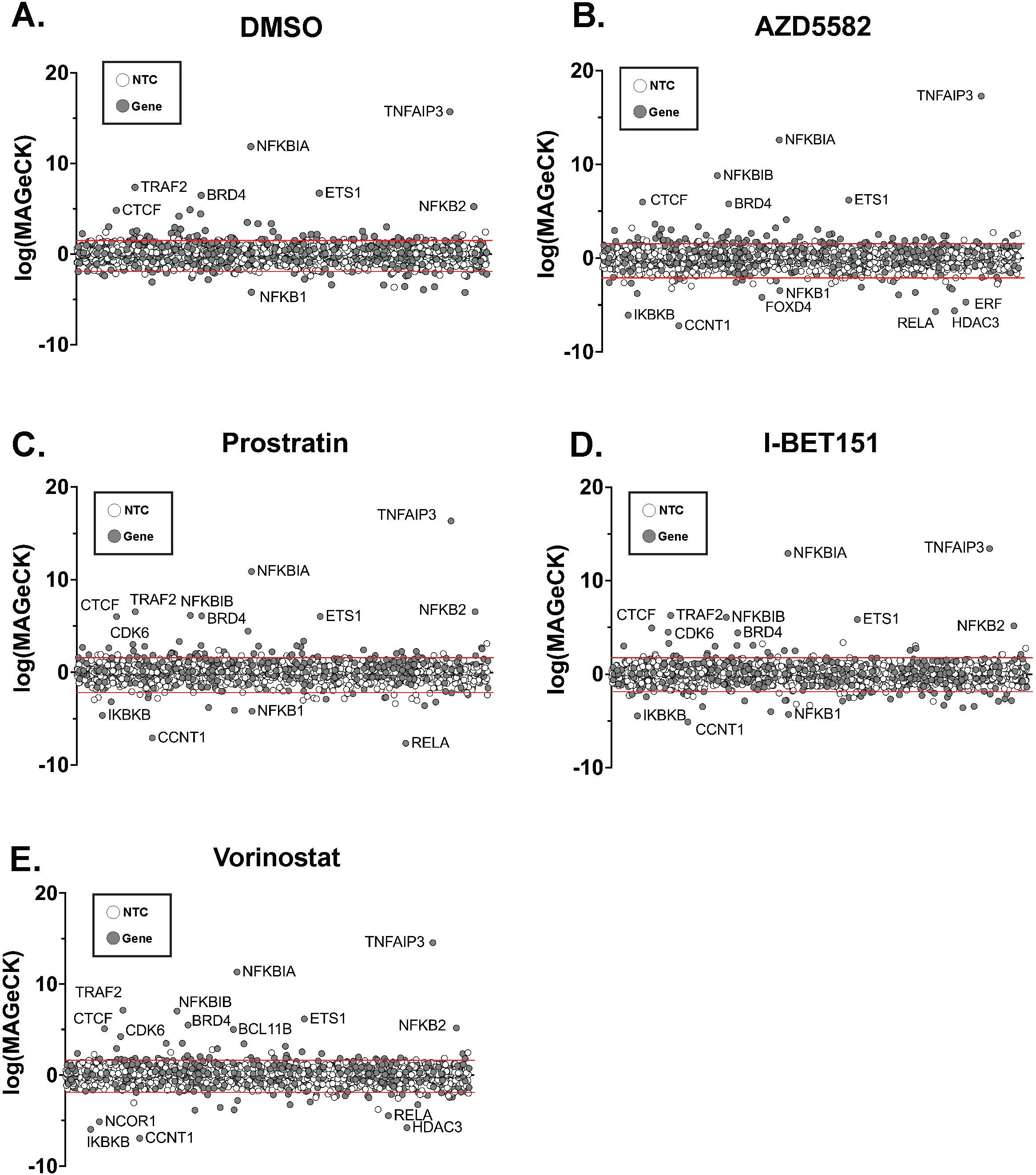
CRISPR Screen of TxLatent library. The Latency HIV-CRISPR Screen is displayed for J-Lat 10.6 cells transduced with TxLatent library and subsequently treated with DMSO (A), AZD5582 (10nM) (B), prostratin (75nM) (C), iBET151 (75nM) (D), or vorinostat (500nM) (E). Genes in the TxLatent library are randomly displayed on the x-axis, in the same order across each treatment. The -Log(MAGeCK Score) is plotted for the enriched genes and the Log(MAGeCK score) is plotted for depleted genes, and combined together to display on a single graph for each treatment. A list of “synthetic” NTCs (synNTCs) was created by randomly grouping sets of 8 NTCs and used to generate a cutoff for gene hits that were enriched or depleted. SynNTCs are shown with white circles and genes targeted by gRNAs in the TxLatent library are shown with gray circles. The red lines on each graph represent the two standard deviations of mean of the synNTCs: DMSO, 1.52 and -1.91; AZD5582, 1.53 and -2.10; iBET151, 1.75 and -1.87; prostratin, 1.58 and -2.19; and vorinostat, 1.61 and -1.9.

We then focused on genes that were positively enriched in this screen (e.g. genes whose knockout caused HIV to release from latency) since these could represent targets for potential latency reversing agents in a “shock and kill” strategy. We observed a substantial number of gene hits enriched in the DMSO (56 genes), AZD5582 (69 genes), iBET151 (28 genes), prostratin (55 genes), and vorinostat (48 genes) conditions. Among the most highly enriched hits in cells treated with the DMSO control (**Figure 2A**), as well as with each LRA treatment, were several negative regulators of the NF-κB signaling. Notably, the top enriched gene for each treatment condition was *TNFAIP3* (**Figure 2A-E**). *TNFAIP3* encodes the Tumor Necrosis Alpha Induced Protein 3, also known as A20, and has been shown to be a key negative regulator of NF-κB signaling by several groups (15, 16). Moreover, *NFKBIA* and *NFKBIB* (NF-κB inhibitors Alpha and Beta, respectively) targeting sgRNAs are consistently positively enriched in secreted viruses with each LRA treatment (**Figures 2B-E**). These genes encode proteins that negatively regulate NF-κB signaling by sequestering NF-κB in the cytoplasm (17). Interestingly, three genes encoding members of the cohesin complex were identified in our analysis as playing a role in repression of HIV: *CTCF*, *Rad21* and *SMC3* (**Figure 2**, **Table S2**). We observed that these sgRNAs targeting these genes were enriched in the screen in J-Lat 10.6 cells as well as in several of the J-Lat 5A8 conditions (**Figure 2**, **Table S2**). The proteins encoded by these genes form a complex that is responsible for mediating long range DNA interactions (18, 19), potentially suggesting a role for this complex or three-dimensional chromatin structure in HIV transcriptional regulation.

We also observed several novel hits, including the transcriptional regulator *ETS1* and the cyclin-dependent kinase *CDK6*. *CDK6* targeting gRNAs were also enriched in the supernatant for prostratin, iBET151 and vorinostat treatments, indicating a role in HIV expression (**Figure 2C-E**). CDK6 has not previously been implicated as a latency regulating factor, though other CDKs such as CDK9 and CDK7 have well described roles in the regulation of RNApol2 (20). *ETS1* targeting sgRNAs were enriched in DMSO exposed cells and for all LRA conditions indicating a general role in repression of HIV. Curiously, previous work has identified ETS1 as a potential regulator of HIV, but these reports suggested that ETS1 cooperates with NF-κB to enhance transcriptional activity of HIV enhancers at the LTR (21, 22). Nevertheless, in this screen, we observe an apparent negative regulatory role for ETS1 in HIV expression.

### Primary CD4 T cell CRISPR validation for novel factors in HIV-1 latency

From the candidate HIV repressing genes identified by the lentiviral CRISPR screen in the Jurkat cell line model, we selected 10 significantly enriched host cellular genes/transcription factors, including AAVS1, CDK6, ETS1 FOXE3, SAMD12, SMC3, TNFAIP3, ZIC5 and ZNF740, for further validation in a primary CD4 T cell latency model (**Figure 3A**). In this model, primary CD4 T cells from seronegative donors are activated via their TCR, then infected with a GFP-expressing reporter strain of HIV (HIV-GFP) (23, 24) followed by enrichment of infected cells at 3 days post infection. After infection, the cells were maintained in culture with IL-2 for up to three weeks, during which time viral gene expression progressively declines as the cells revert to a resting state, and a population of latently infected (GFP-) cells emerges. Three days post enrichment, cells were nucleofected with Cas9/sgRNA ribonucleoprotein (RNP) complexes designed against each target. RNPs with scrambled non-targeting sgRNA or Tat-targeting sgRNAs were nucleofected in parallel as negative and positive controls respectively (**Figure 3A**). Viral gene expression was then measured by flow cytometry for GFP at 2-week post nucleofection to determine the effect on HIV latency (**Figure 3B**). As expected, CRISPR targeting of Tat led to a potent reduction in viral expression. Interestingly, viral expression was significantly elevated for cells in which ETS1 was targeted (p<0.0001) compared to NT control (**Figure 3B, 3C**). These data indicate that as the infected CD4 T cells return to a resting state, ETS1 plays an important role in establishing repression of latent HIV.

**Figure 3.**
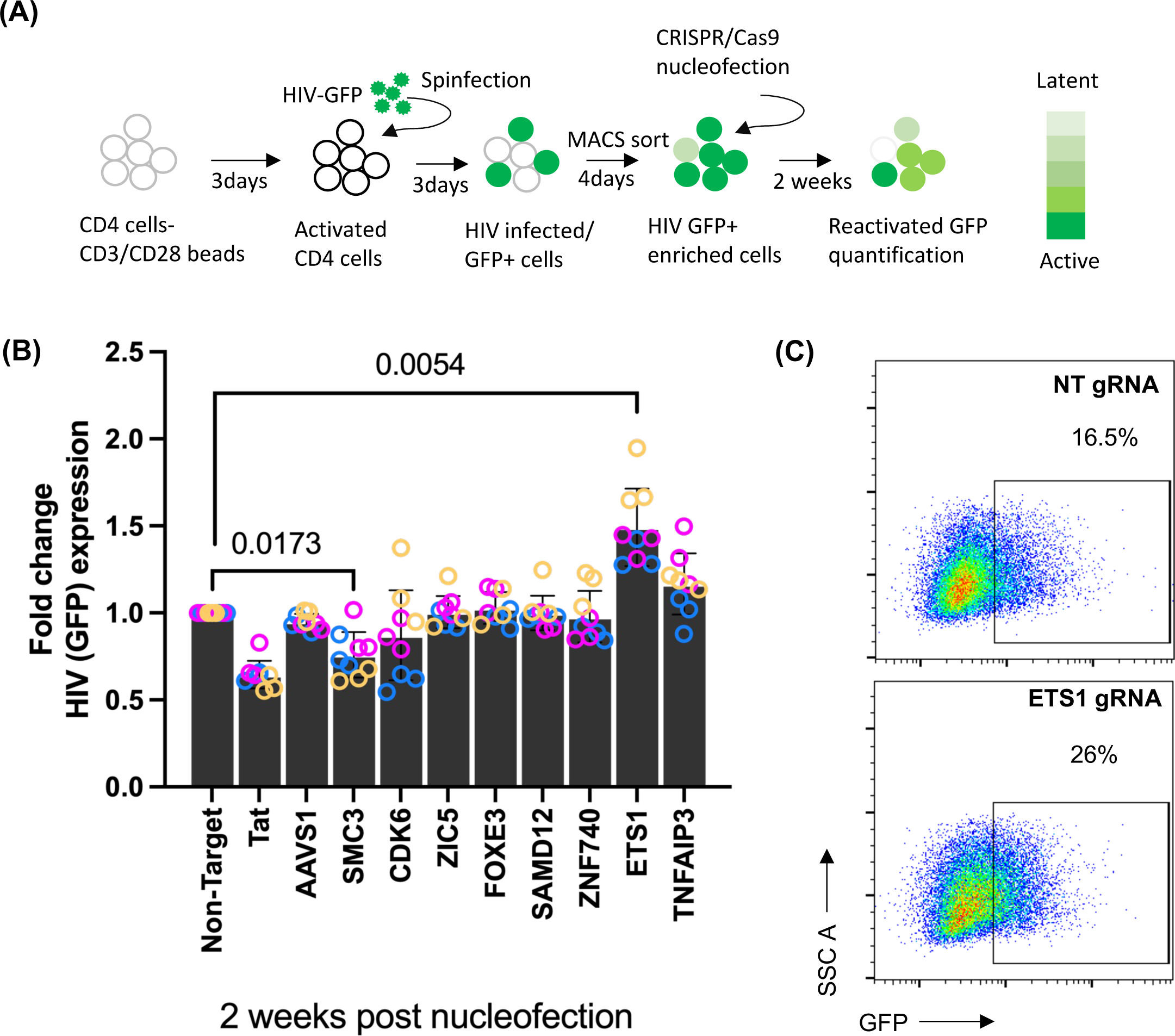
CRISPR validation of targets in HIV infected primary CD4 T cells. **(A)** Schematic overview of experimental design for CRISPR-Cas9 targeting of host genes in HIV-GFP infected primary CD4 T cells. Activated CD4 T cells were infected with HIV-GFP (23). At 3 days post infection, infected (GFP+) cells were enriched by magnetic sorting and cultured for 4 days before nucleofection of Cas9/sgRNA complexes targeting AAVS1, SMC3, CDK6, ZIC5, FOXE3, SAMD12, ZNF740, ETS1, and TNFAIP3 or a non-targeting sgRNA (NT). Pools of 3 different gRNAs were used for each target. HIV infected cells are denoted in green color. Differential intensity of green color represents latent and active infection. **(B)** Relative level of HIV expression measured as GFP using flow cytometry. GFP expression was measured at two weeks post nucleofection (wpN). Data are displayed as GFP expression in fold change normalized to cells nucleofected with NT gRNA. Each condition represents three biological replicates/donors (Yellow, Blue and Purple) and three technical replicates for each donor. **(C)** Dot plots representing viral GFP expression at two weeks post knockout. Statistical analysis was conducted for cumulative data. Error bars represent standard deviations, and P values displayed were determined by two-way ANOVA Tukey’s multiple comparisons test. WT, wildtype; NT, non-target; CRISPR, clustered regularly interspaced short palindromic repeats; Cas9, CRISPR-associated protein 9.

### ETS1 promotes HIV expression in actively infected cells and inhibits HIV expression in latently infected cells

ETS1 has previously been reported to positively regulate HIV expression in complexes with other cellular transcription factors, like NF-κB, NFAT, and USF-1 (25, 26). An ETS1 binding site has been identified in the LTR distal region, and other ETS family members have also been shown to bind the LTR (27). To reconcile these previous observations with our finding that ETS1 participates in repressing latent HIV, we examined the impact of ETS1 knockout at different timepoints post infection as well as in the presence of latency reversing agents (LRAs). In our latency model, T cells are first activated through TCR mediated signaling, and slowly return to a resting state over two weeks. RNPs with sgRNAs targeting ETS1, the HIV Tat transcriptional regulator or a non-targeting control were electroporated into recently activated infected primary cells to examine the impact on HIV-1. Western blot analysis of nucleofected cells confirmed nearly complete depletion of ETS1 at 1- and 2-week post nucleofection (**Figure 4A**). Strikingly, when we examined an early timepoint (1wpi) post infection when the cells are still in an activated state and majority of infected cells are actively infected, we observed that ETS1 depleted cells had reduced HIV expression compared to non-targeting controls (**Figure 4B and C)**, while at later timepoints (2wpi) we observed an increase in HIV expression for ETS1 depleted cells, relative to controls (**Figure 4D and E**), consistent with our previous observation (**Figure 3B**). Thus, ETS1 appears to play distinct roles at different times post T cell activation – a positive role at earlier times while the cells are still in a recently activated state, but a repressive role at later timepoints when the cells have returned to a resting state.

**Figure 4.**
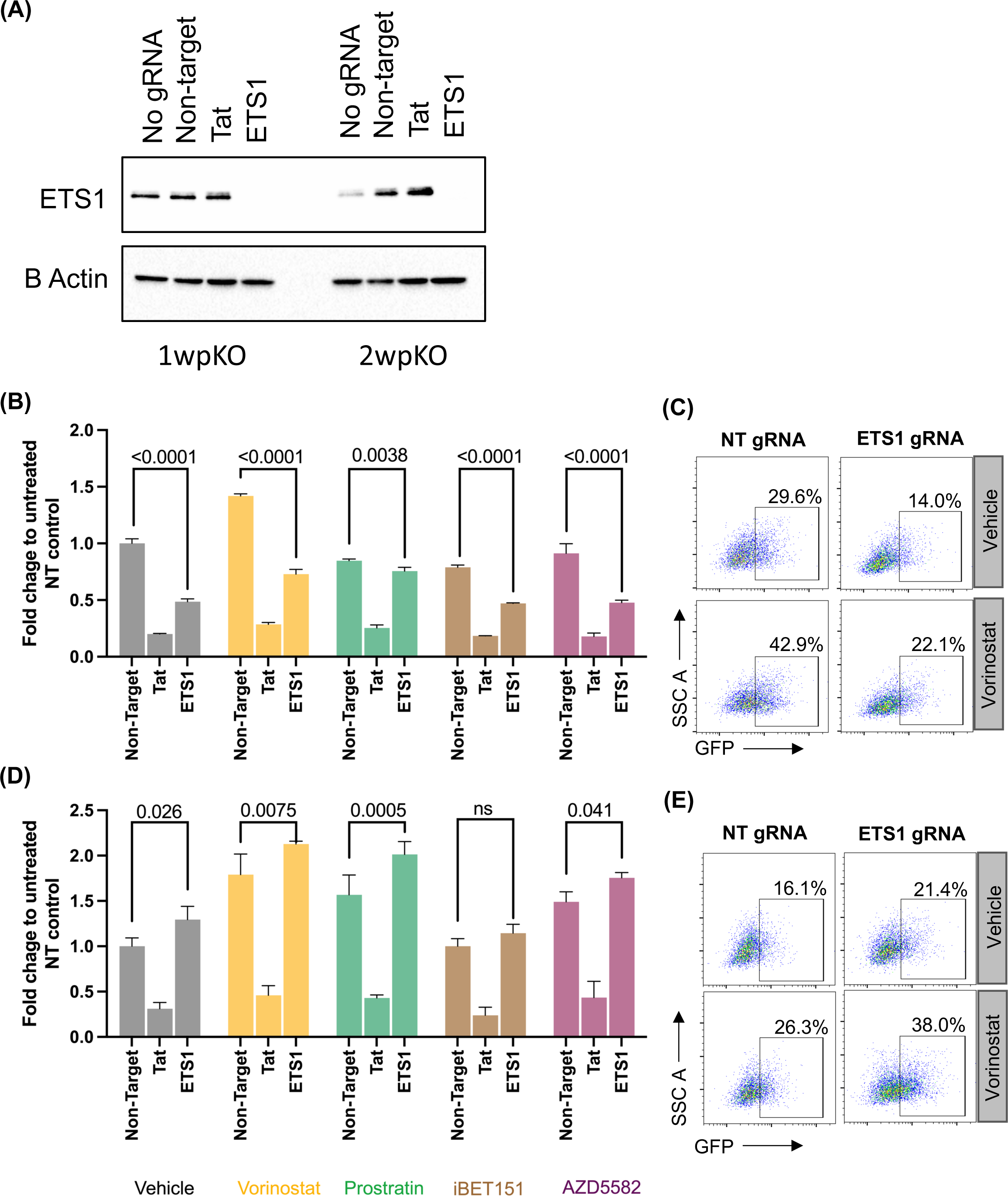
ETS1 knockout in primary CD4 T cells reveals positive and negative roles in HIV expression. Primary CD4 T cells were activated, infected with HIV-GFP, enriched and nucleofected as in Figure 3. **(A)** Western blot of the ETS1-targeting or a control non-targeting (NT) Cas9/gRNA nucleofected primary cells for beta actin and ETS1 expression at one and two weeks post nucleofection (pKO). **(B-E)** Flow cytometry of infected cells without stimulation or after 24 h of stimulation with four different LRAs (vorinostat, prostratin, iBET151, and AZD5582 at one **(B)** and two **(D)** week post nucleofection. **(C and E)** Dot plots representing viral GFP expression at one and two weeks post knockout respectively. Data are represented as fold change in percent GFP expression, normalized to cells nucleofected with NT gRNA. Error bars represent standard deviation, and P values displayed were determined by two-way ANOVA Tukey’s multiple comparisons test.

We also evaluated the role of ETS1 in latently infected primary CD4 T cells in response to different LRAs that we used to screen the different target in Jurkat cell line models. Specifically, we stimulated the infected cells with a panel of LRAs with distinct mechanisms of action: Vorinostat (HDACi), Prostratin (PKCa), iBET151 (BETi), and AZD5582 (non-canonical NF-κB agonist) at 500nM for 24h. At 1wpi this stimulation caused a significant increase in the percentage of GFP+ cells in the cell population for Vorinostat but caused only a small response to prostratin and no apparent response to iBET151 or AZD5582. This increase also occurred in ETS1 depleted cells. However, at two weeks post nucleofection, depletion of ETS1 caused augmented viral gene expression at base line as expected in all drug conditions. At 2wpi, both ETS1 depleted cells and control cells also responded to Vorinostat, Prostratin, and to AZD5582 but not iBET151. These data indicate that ETS1 is important cellular host factor for both expression of HIV in active cells and establishment of latent HIV infection in resting cells, but that ETS1 depletion does not noticeably change the sensitivity of latent proviruses to benchmark LRAs.

### ETS1 is required for repression of latent HIV in cells from ART-suppressed people with HIV

Having found that ETS1 contributes to suppressing HIV expression in resting CD4 T cells that had been infected ex vivo, we next examined whether ETS1 is required for maintaining HIV latency in the clinical reservoir. To investigate this, CD4 T cells were isolated from three ART suppressed people with HIV (PWH). Isolated total CD4 T cells were rested for 24 h followed by nucleofection with non-targeting or ETS1 targeting gRNAs. A schematic of CRISPR/Cas9 based nucleofection in CD4 T cells from PWH is depicted in **Figure 5A**. At 3 days post nucleofection, complete depletion of ETS1 was seen in cells electroporated with ETS1-targeting guide RNAs by western blot (**Figures 5B, S3A**). To determine the effect of ETS1 depletion on HIV expression, we then carried out quantitative PCR for Gag unspliced viral RNA for each sample. Notably, we observed robust upregulation of HIV-Gag mRNA expression within the ETS1 depleted cells, with elevated relative HIV-specific mRNA expression levels ranging at 8.2, 2.3, and 2-fold change (**Figures 5C, S3B**) compared to non-targeting (NT) guide RNAs with statistical significance (p<0.0001, 0.0001, and 0.0015, respectively; 2-way ANOVA Tukey’s multiple comparisons test). These findings confirm that ETS1 is required for the maintenance of latent HIV infection in resting cells *in vivo*.

**Figure 5.**
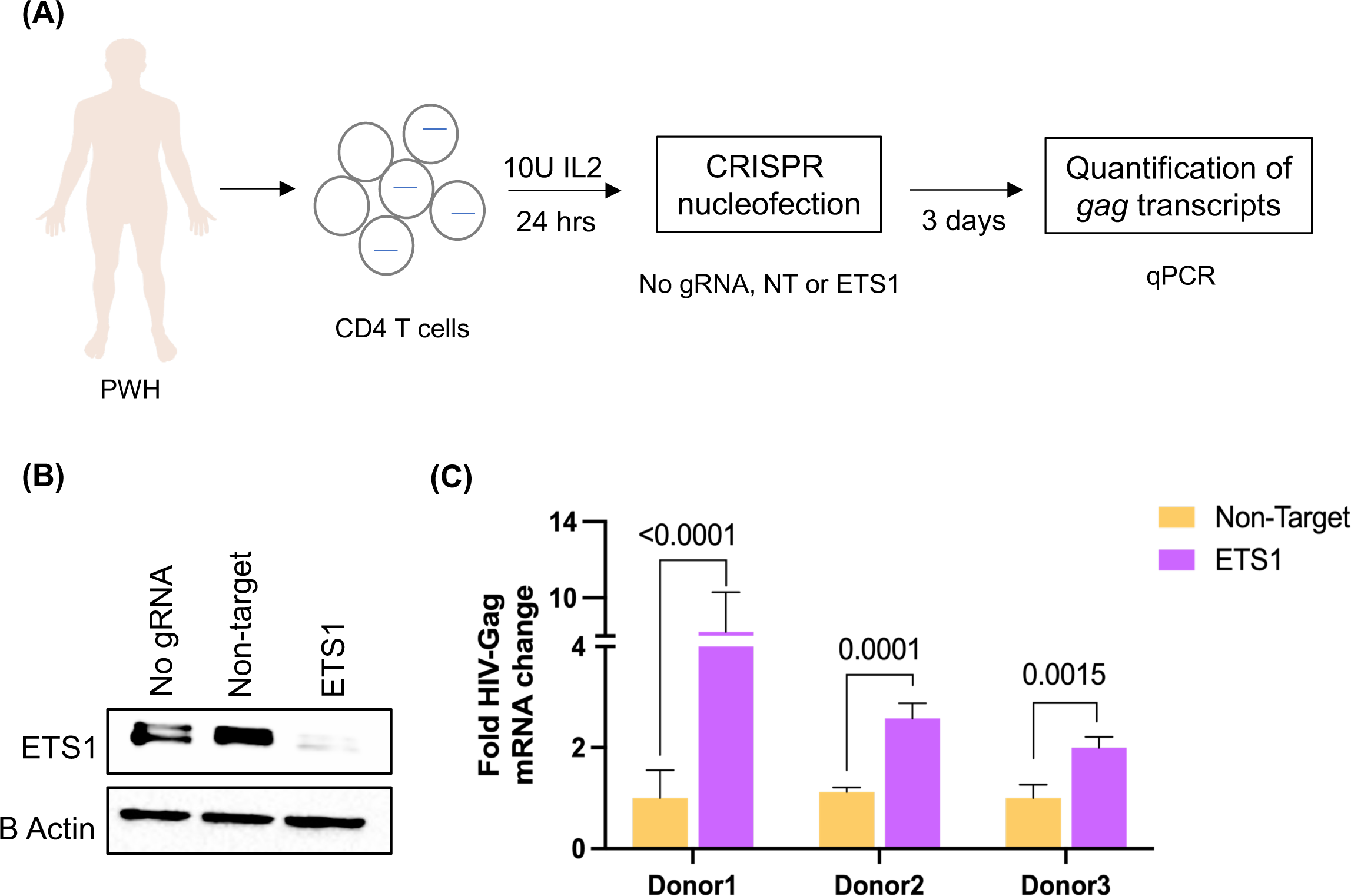
ETS1 knockout reactivates HIV transcription in CD4 T cells from people with HIV (PWH) on antiretroviral therapy (ART) *ex vivo*. **(A)** Schematic of *ex vivo* CRISPR nucleofection experiment in resting CD4 T cells isolated from PWH on ART. CRISPR nucleofection of ETS1-targeting or non-targeting (NT) Cas9/gRNAs was performed after a 24 hr culture of PWH CD4 T cells followed by quantification of Gag RNA expression at 3 days post nucleofection by quantitative PCR. **(B)** Depletion of ETS1 in nucleofected resting CD4 T cells was analyzed by western blot. **(C)** Relative abundance of Gag viral RNA expression in CD4 T cells nucleofected with ETS1-targeting or NT control Cas9/gRNA complexes. Gag viral RNA expression was normalized to cells nucleofected with no gRNA and presented as fold change. Experiment was conducted in three biological replicates/donors with three technical replicates for each condition. Error bars represent standard deviation, and P values displayed were determined by two-way ANOVA Tukey’s multiple comparisons test.

### Transcriptomic profiling of ETS1 depleted CD4 T cells identifies several known and novel candidate regulators of HIV gene expression in cells ex vivo

Although our data demonstrate that ETS1 can play a suppressive role in HIV expression and helps to maintain silencing of the HIV reservoir, the mechanisms by which ETS1 contributes to HIV latency and reactivation in PWH are not completely characterized. To further understand the role of ETS1 in regulation of HIV expression, we analyzed the transcriptome of CD4 T cells from PWH with and without knockout of ETS1. RNA from the ETS1-targeting or non-targeting sgRNA nucleofected PWH donors (**Figure 5A**) were analyzed by RNA sequencing (RNAseq). Principal component analysis (PCA) was used to determine the overall impact of ETS1 knockout on gene expression patterns across the conditions. Importantly, the samples clustered by knockout condition, indicating a consistent transcriptomic impact of ETS1 knockout across the donors (**Figure 6A**). We identified several differentially expressed genes (DEGs) between the two conditions, 178 genes total (padj<0.05), with 86 genes upregulated and 92 downregulated (**Figure 6B, Table S3**).

**Figure 6.**
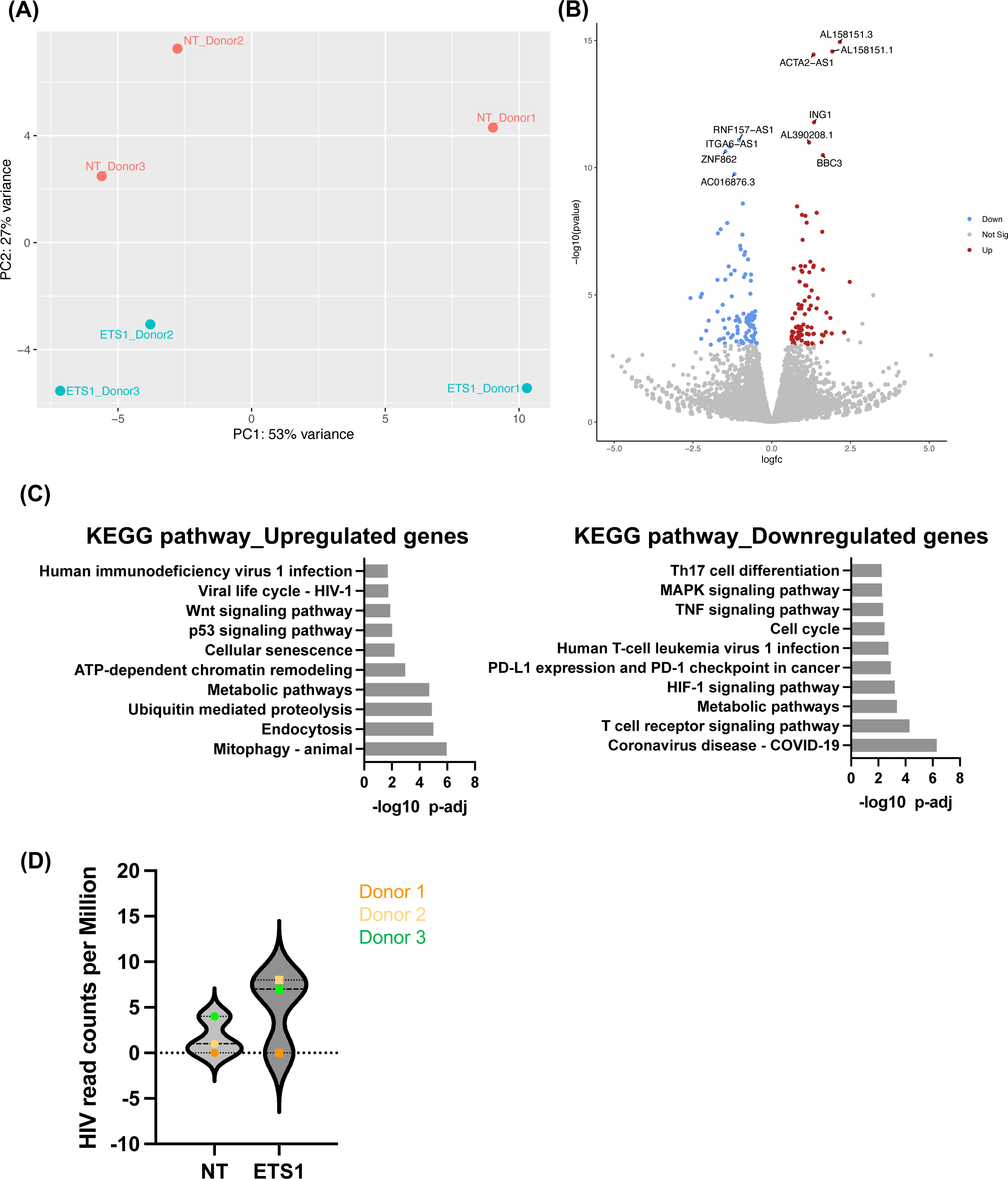
Analysis of differential gene expression in CD4 T cells from PWH on ART after depletion of ETS1. CD4 T cells from three PWH on ART were nucleofected with Cas9/gRNA complexes targeting ETS1 or non-targeting control (NT) as described in Figure 5, followed by bulk RNAseq. **(A)** Principal component analysis (PCA) plot of samples based on gene expression data for each sample. ETS1-targeted samples shown by blue dots, non-targeting gRNA samples by red dots. Each data point represents an individual sample. The y-axis and x-axis represent the first and second principal components, respectively. **(B)** Volcano plot of overall differentially expressed genes (DEGs) between HIV infected CD4 T cells nucleofected with ETS1 targeting sgRNA/Ca9 complexes vs non-targeting. **(C)** Database for Annotation, Visualization and Integrated Discovery (DAVID) analysis of upregulated and downregulated genes of the RNA-sequencing analysis of ETS1 vs control cells. The 10 most significant KEGG pathways are shown. **(D)** Number of unique HIV-mapping reads identified in each sample after alignment to an HXB2 reference genome.

Interestingly ETS1 was not identified as a DEG by this analysis, despite being clearly depleted by western blot, prompting us to examine ETS1 mapping reads in the dataset in more detail. For ETS1 targeting, three ETS1-specific sgRNAs were used in combination (sgRNA1, sgRNA2, and sgRNA3). When we examined read coverage between the recognition sites for sgRNA1 and sgRNA2 we observed a clear drop in read coverage (**Figure S4A**), indicating that the sgRNAs caused a disruption to the ETS1 gene within this region. (28-30) Thus, although the overall number of RNAseq reads within ETS1 did not change with sgRNA targeting, the gene is clearly affected by the sgRNAs, leading to reduced ETS1 protein expression.

We also aligned the RNAseq dataset to the HXB2 reference HIV genome to examine the abundance of HIV mapping reads. As expected, HIV mapping reads were rare and coverage across the HIV genome was extremely sparse. For one donor, no HIV reads were observed, but HIV RNA was detected for the other two donors. We observed 1 and 4 unique HIV reads for NT samples and 8 and 7 unique HIV reads for ETS1 depleted cells respectively for Donor 2 and 3 (**Figure 6D, Figure S4B**), consistent with the hypothesis that ETS1 knockout reactivates HIV expression.

To understand the overall impact of ETS1 knockout on CD4 T cells from PWH, we next examined the pathways represented within the set of DEGs. Interestingly, some of the top upregulated transcripts are involved in mitophagy, endocytosis, metabolic pathways, ATP-dependent chromatin modeling (**Figure 6C, Left panel**). ATP-dependent chromatin-remodeling complexes play an important role in HIV transcription as these complexes are recruited by HIV Tat to the long terminal repeat (LTR) to facilitate active transcription (31). Notably, ETS1 depletion also upregulated genes involved in the Wnt signaling pathway, HIV-1 viral life cycle, and HIV-1 infection pathways (**Figure 6C**). Downregulated pathways included COVID-19, T cell receptor signaling pathway, metabolic pathway, and HIF-1 signaling pathway (**Figure 6C Right panel**). HIF-1 signaling pathway is involved in viral replication and inflammation through extracellular vesicles (32). ETS1 depletion also downregulated transcripts involved in MAPK signaling pathway, TNF signaling pathway, and Th17 cell differentiation that are important in several cellular responses including expression of several inflammatory cytokines.

Expression of the ING1 (Inhibitor Of Growth Family Member 1) was upregulated in response to ETS1 depletion. ING1 is a nuclear protein involved in Chromatin Regulation that physically interacts with p53/TP53 in the negative regulatory pathway of cell growth by modulating p53-dependent transcriptional activation. Transcriptional activators KLF7 and FOXH were also upregulated in response to ETS1 depletion. Additionally, we observed upregulation of the cytokine receptor IFNGR2 (Interferon Gamma Receptor 2) that is critical for innate and adaptive immunity against viral, some bacterial and protozoan infections in response to ETS1 depletion. We also observed increased expression of the interferon-stimulated gene (ISG), TRIM5, a cytoplasmic antiretroviral effector (33) involved in inflammation (34) and autophagy (35). By contrast, IL27RA, a critical gene for cell-mediated immunity, and TNFRSF25 - a mediator of NF-κB activation and apoptosis, were downregulated. Several novel lncRNAs, PVT1, DM1-AS, ZNF337-AS1 were also significantly expressed in response to ETS1 depletion.

We observed several known and novel candidate regulators of HIV gene expression in the RNAseq dataset. In particular, we observed that expression of the long non-coding RNA MALAT1 (metastasis-associated lung adenocarcinoma transcript 1) was upregulated in ETS1 depleted cells. MALAT1 is a well-known long non-coding RNA, that has been previously reported to promote HIV transcription and infection (36, 37). MALAT1 influences gene expression by several mechanisms, including regulating the recruitment of specific chromatin modifiers such as the Polycomb Repressive Complex 2 (PRC2). Interestingly, one of the top upregulated transcripts in ETS1 depleted cells was a long noncoding gene, AL158151.3, that has recently been reported as being associated with HIV control (38). Overall, these data demonstrate that ETS1 depletion influences several cellular genes, including some that could have a direct or indirect impact on HIV expression.

### Reactivation of HIV following ETS1 depletion is partially dependent on MALAT1

Having observed a potential impact of ETS1 on MALAT1 expression in HIV-1 infected cells, we next further investigated the impact of ETS1 on MALAT1 and HIV expression. Using our primary CD4 T cell model of HIV latency, we nucleofected cells from three independent donors with RNPs containing sgRNAs targeting ETS1, MALAT1, or both ETS1 and MALAT1 in combination. Two weeks post nucleofection of HIV-GFP infected CD4 T cells, the protein level of ETS1 was quantified by immunoblotting (**Figure 7A**). Additionally, we measured the abundance of MALAT1 and HIV-1 Gag RNA in the cells by qRT-PCR (**Figure 7B and 7C**), and viral protein expression by flow cytometry (**Figure 7D**). As expected, ETS1 was strongly depleted by ETS1 targeting sgRNAs, but not by control sgRNAs (**Figure 7A**). MALAT1 expression was also significantly depleted in all three donors for cells with sgRNAs targeting MALAT1 (**Figure 7B**). We observed that depletion of ETS1 increased MALAT1 RNA expression from latently infected cells (**Figure 7B, 7C**), although the magnitude of the change was variable from donor to donor. ETS1 depletion also caused significant upregulation of HIV RNA in two of the three donors, while a trend towards an increase was observed in the third donor. Notably, the magnitude of HIV RNA upregulation followed the same pattern as the increase in MALAT1 expression. For HIV protein expression, two of the three donors demonstrated a significant increase in GFP after depletion of ETS1, while one donor did not (**Figure 7D**). Depletion of MALAT1 alone had no effect on HIV expression. In contrast, depletion of MALAT1 in latently infected CD4+ T cells in combination with ETS1 knockout decreased HIV-1 reactivation, as determined by quantification of cell associated Gag and GFP expression (**Figure 7C and 7D**) although this effect was more pronounce for some donors than others. One donor (donor 2) did not exhibit any reactivation of HIV with ETS1 depletion at the protein level (**Figure 7D**), but exhibited an impact on HIV RNA level (**Figure 7C**), suggesting that for some donors, additional posttranscriptional blocks continue to inhibit HIV protein expression, even in the presence of ETS1 knockout. These results indicate that ETS1 represses HIV expression at the transcriptional level in latently infected CD4 T cells, and that this effect is partly dependent on ETS1-mediated repression of MALAT1 expression, although donor-to-donor variation exists for this interaction.

**Figure 7.**
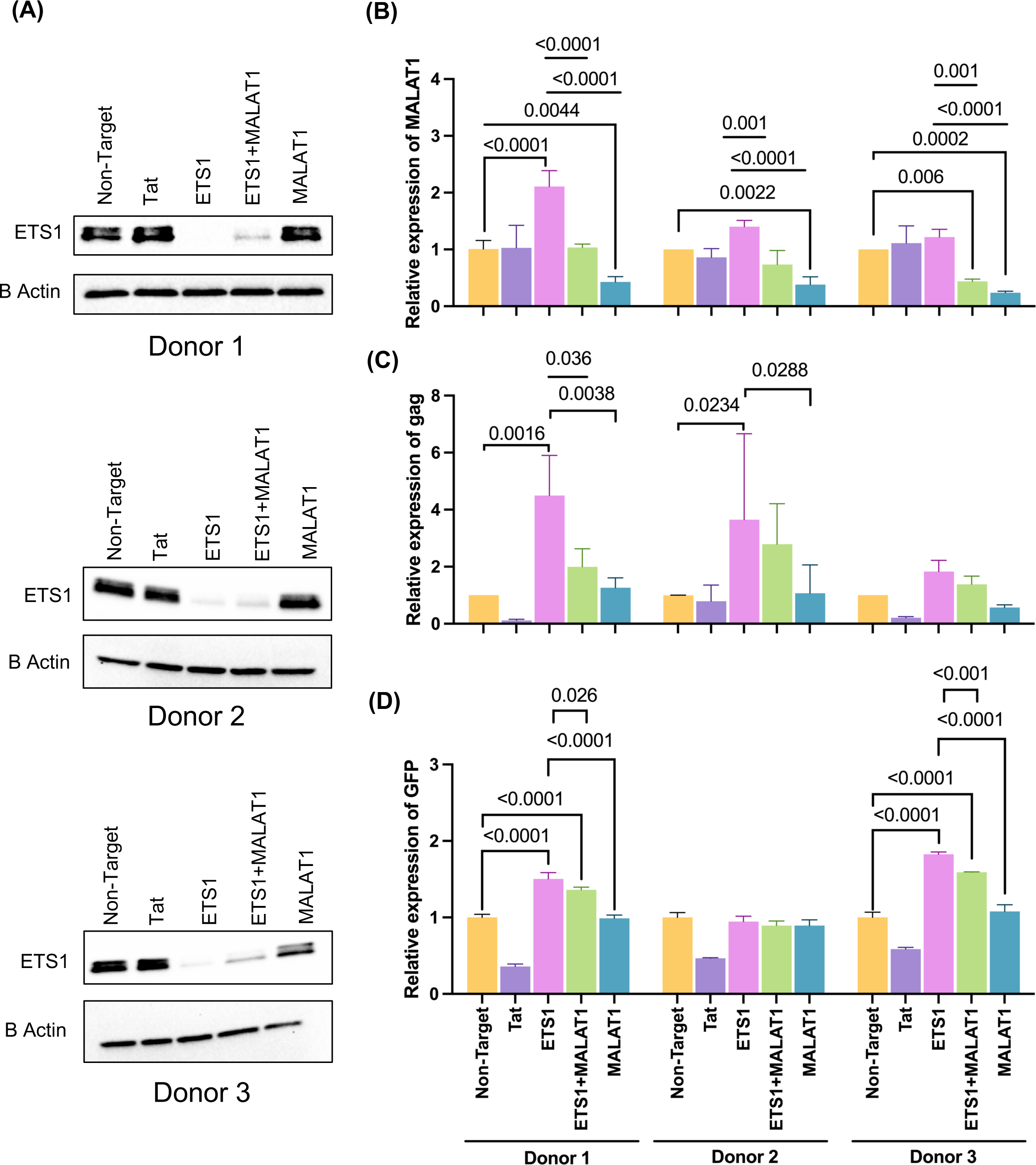
Reactivation of HIV following ETS1 depletion is partially dependent on MALAT1. *In vitro* generated primary latent CD4 cells were nucleofected with Cas9/gRNA ribonucleoprotein complexes targeting ETS1, MALAT1, or a combination of ETS1 and MALAT1 as described in Figure 5, followed by immunoblotting, flow cytometry and RT-PCR. (**A**) Depletion of ETS1 in nucleofected CD4 T cells was analyzed by western blot. (**B-C**) Relative abundance of MALAT1 expression (**B**), and Gag RNA (**C**) in CD4 T cells nucleofected with Cas9/sgRNAs targeting ETS1, MALAT1, or a combination of ETS1 and MALAT1. (**D**) Flow cytometry of infected cells at two-week post nucleofection. MALAT1 and Gag viral RNA expression was normalized to cells nucleofected with NT control. Experiment was conducted in three biological replicates/donors with three technical replicates for each condition. Error bars represent standard deviations, and P values displayed were determined by two-way ANOVA Tukey’s multiple comparisons test.

## Discussion

Complex interplay between numerous viral and cellular transcription factors determines the molecular regulation of HIV-1 transcription, and the HIV-1 promoter contains binding sites for several cellular TFs with either activating and repressive functions (5). For example, transcription activators that bind to the HIV-1 5′ LTR include Sp1, NF-κB, the AP-1 complex family, NFAT, and the long non-coding RNA, MALAT1 (36, 39-42). Transcriptional silencers also bind the HIV-1 5′LTR, including NELF, YY1, ZNF304, and AP4 (43-46). However, the overall nature of how these factors interact to determine HIV expression is still not well understood and it is likely that additional host factors that regulate HIV exist. Thus, a deeper understanding of the host cell activators and repressors that HIV-1 relies on for both latency or viral gene expression will be required for the effective development and implementation of curative approaches that target HIV expression.

To address this gap, we have employed single-cell technologies (integrated single cell RNA and ATAC sequencing) (6) together with CRISPR-Cas9 lentiviral genomic screening (11) in J-Lat 10.6 and J-Lat 5A8 cells to identify a set of novel regulators of HIV expression. In this present study, hundreds of cellular transcripts that were correlated with HIV transcription identified in our previous study (6) were tested for a role in HIV latency by a new latency CRISPR screening strategy called lentiviral CRISPR screening. This approach revealed several novel genes involved in latency, including ETS1, TNFAIP3, ZNF740, SAMD12, FOXE3, ZIC5, SMC3 and CDK6. After carrying out further validation in primary CD4 T cells *in vitro* with a small number of selected targets, we confirmed that ETS1 represses HIV expression and maintains latency in primary CD4 T cells. We also identified a member of the cyclin-dependent kinase family (CDK6) and a cohesin complex subunit, structural maintenance of chromosomes (SMC3) significantly impact HIV expression in primary cells. SMC3 was previously reported to involved in transcriptional regulation and epigenetic processes by binding to HIV-1 Tat (47). Shimura et al in 2011 discovered that expression of the HIV-1 protein Vpr decreases the recruitment of SMC3 to chromatin (48). Several CDK family members have previously been documented for their association with HIV infection and replication (49). For example, CDK1, CDK2, and CDK6 phosphorylate SAMHD1 and inactivate its antiviral function (49, 50). CDK7, CDK11, and CDK13 were reported to be involved in RNAP II initiation of transcription at the HIV-1 provirus, HIV RNA 3’ end processing, and HIV-1 RNA splicing respectively. Recent studies have discovered that CDK8 and CDK19 (51), and CDK9 (52) induce the expression of latent HIV provirus. However, in the present study, we prioritized ETS1 for additional investigation, as it appears to exhibit a repressive role in HIV expression in latently infected cells, thus representing a novel target for LRAs.

ETS1 is the prototype of the ETS family of transcription activators and plays an important role in host transcriptional response and in regulation of immune cell function (21), T cell activation and cytokine expression (53), and viral infection (54). ETS1 has been shown to function either as transcriptional activator or repressors of numerous genes (55), and is involved in stem cell development, cell senescence and death, and tumorigenesis (56). Previous work using overexpression to examine the impact of ETS1 on HIV promoter activity in transformed cell lines demonstrated that ETS1 can promote HIV expression, and that an ETS1 binding site exists in the HIV LTR. The divergent behavior of ETS1 with respect to HIV expression at different times could result from different complexes or binding partners of ETS1. Curiously, it has been previously shown that the ETS1 binding site in HIV is not required for activation of the LTR by a truncated version of ETS1, suggesting a possible indirect effect of ETS1 on HIV (26). In addition to validation studies in an *in vitro* latency model, we also performed ETS1 depletion *ex vivo* in resting CD4 cells isolated from ART suppressed person with HIV and observed viral reactivation. This discovery clearly demonstrates that ETS1 can play a repressive function in the host cells of the HIV reservoir.

We hypothesize that the depletion of ETS1 affects HIV indirectly. Consistent with this hypothesis, RNAseq profiling of ETS1 depleted CD4 T cells identified HIV regulating genes as being significantly enriched within the set of differentially expressed genes. Previous studies have also indicated diverse functions for ETS1 in T cells. ETS1 is essential for the activation of T cells and the production of IFN-γ (57, 58). In a recent study of Helicobacter pylori-associated gastritis, ETS1 expression was shown to be induced by infection and to activate proinflammatory gene expression (59). ETS1 has also been reported to regulate malignant tumor cell invasion (60), restricts virus replication and is induced by poxvirus infection (54). It should be noted that inhibitors for ETS1 are not yet available commercially or clinically. However, recent studies described novel ETS1 inhibitors and their role in antitumor activity against hepatitis B/C virus (HBV/HCV) induced hepatocellular carcinoma (60), lymphoma (61) and vessel regression (62).

Interestingly, in our RNA-Seq data, we observed that ETS1 depletion in resting CD4 T cells upregulates MALAT1, a known positive regulator of HIV expression. This observation suggests a potential regulatory relationship between ETS1 and MALAT1 in the context of HIV infection. Association of MALAT1 expression in HIV-1 transcription and reactivation is reported in several studies (36, 63). MALAT1 in HIV infection interacts with PRC2 responsible for the methylation (di- and tri-) of Lys 27 of histone H3. Facilitation of MALAT1-PRC2 complex protects the HIV-1 LTR from PRC2-mediated epigenetic silencing, which resulted in decreased H3K27 trimethylation on LTR, and hence HIV transcription (63, 64). Additionally, depletion of ETS1 caused the upregulation of cellular genes that are involved in ATP dependent chromatin remodeling. Chromatin remodelers play a critical role in regulating multiple cellular processes, such as DNA repair and gene expression, creating nucleosome-depleted regions by promoting the exchange of histones (65). Such chromatin remodeling complexes are also involved in regulating the chromatin architecture at the HIV LTR thereby regulating LTR transcription (66). Alterations to chromatin remodeling may thus contribute to the latency regulating activity of ETS1.

Since the observations from the RNAseq data showed upregulated MALAT1 expression in ETS1 depleted resting cells from PWH on ART, we employed the same CRISPR knockout approach in *in vitro* primary latent HIV cell model to address the mechanistic role of ETS1 and MALAT1 in HIV-1 latency. Our data from the three independent donors demonstrates that ETS1 knockout results in a trend for increased HIV-1 and MALAT1 RNA in infected cells, although the findings were somewhat variable between donors. Combined depletion of MALAT1 with ETS1 indicated that this attenuated the upregulation of viral gene expression induced by ETS1 knockout, suggesting that the impact of ETS1 knockout on HIV-1 is partly dependent on MALAT1 upregulation. Indeed, the binding of MALAT1 and ETS1 was reported in a recent tumor study (67). The interplay between MALAT1 and ETS1 has not been previously reported for HIV-1 infection, and we propose that this represents a novel indirect mechanism of HIV regulation by ETS1. Notably, ETS1 is polymorphic in humans with several different alleles being present at high frequency (68). These genetic difference may contribute to the variable impact of ETS1 on HIV and MALAT1 in primary CD4 T cells.

Overall, these data indicate that ETS1 is an important regulator of HIV latency and could represent a useful target for latency reversing strategies. Additional work to reveal the precise mechanisms by which which ETS1 mediates positive and negative regulation of HIV will also help to direct the development of novel LRAs that target ETS1.

## Methods

### Cell lines

The Jurkat T-lymphocyte cell lines J-Lat 10.6 (69) and J-Lat 5A8 (70) were maintained in Gibco RMPI medium (Thermo Fisher Scientific), supplemented with 10% fetal bovine serum, 2 mg/ml l-glutamine, Sodium Pyruvate, penicillin-streptomycin and 10mM HEPES. Cells were cultured at 37 °C with 5% CO2. The J-Lat 10.6 (71) and J-Lat 5A8 (70) cells, are a derivative of Jurkat cells that was infected with a pseudotyped HIV-1 strain, HIV/R7/E−/GFP, have different proviral integration sites, with J-Lat 5A8 having a low multiplicity of infection. The human embryonic kidney HEK293T (ATCC; CRL11268) cell line that was used for transfection was maintained in DMEM complete medium (Gibco), supplemented with 10% fetal bovine serum, and penicillin-streptomycin.

### PWH Participants and Primary CD4 T cell model of HIV latency

Ficoll-Paque (Thermo Fisher Scientific) was used to purify PBMC from leukapheresis donations obtained from durably suppressed, on ART PWH under an approved protocol by the University of North Carolina (UNC) Institutional Review Board. All participants provided informed consent. Total CD4 T cells were isolated from PBMC using EasySep™ Human CD4+ T Cell Isolation Kit (#17952, StemCell Technologies) following the manufacturer’s protocol. CD4 T cells isolated from ART-suppressed people with HIV (PWH) were cultured in RPMI media supplemented with 10 units per mL of IL-2.To generate primary CD4 T cells that were latently infected with HIV, we used a model that we have previously developed (6, 24, 72). Briefly, CD4 T cells from seronegative donors were activated with anti-CD3/CD28 beads (Life Technologies) at 1M cells per mL for 48h, then infected with a replication defective HIV reporter clone, NL4-3-△6-drEGFP-IRES-thy1.2 that had been pseudotyped with the Vesicular Stomatitis Virus G protein (VSV-G) (6, 23). At 48h post infection, infected (GFP+) cells were enriched by magnetic separation of Thy1.2+ cells, then cultured for up to two additional weeks in RPMI media (Thermo Fisher Scientific), supplemented with 10% fetal bovine serum, 2 mg/mL l-glutamine, Sodium Pyruvate, penicillin-streptomycin and 10mM HEPES. Primary CD4+ T cells were additionally supplemented with 20U/mL IL-2 and 4ng/mL IL-7. Cells were cultured at 37 °C with 5% CO2.

### gRNA library generation

Guide RNAs for the 354 HIV correlated genes were generating using GUIDES (73), CHOPCHOP (74) and Synthego gRNA generation algorithms. For each gene, 8 individual gRNAs were generated. The library was supplemented with 150 non-targeting controls (NTCs), to make a total gene library size of 2946 guides (Twist Biosciences, San Francisco, CA). The full list of genes and corresponding guides is available in **Table S4**. DNA sequences corresponding to gRNAs were cloned into the HIV-CRISPR vector as previously described (75). Briefly, BsmBI overhangs were PCR amplified on the guide RNA library and cloned into a BsmBI-digested (New England Biolabs, R0134S) HIV-CRISPR vector by Gibson Assembly (New England Biolabs, E2611S). Representation of guide RNAs in HIV-CRISPR vector was validated by next generation sequencing.

### HIV-CRISPR Library Transduction and Screen

The pool of HIV-CRISPR vectors containing the sgRNA library was transfected into HEK293T cells. Transfection was performed with the library containing HIV-CRISPR vector (667 ng) along with psPax2 (Gagpol, Addgene, 12260, 500 ng) and MD2.G (VSVG, Addgene, 12259), in serum-free DMEM and the TransIT-LT1 reagent (Mirus Bio, MIR2305). Virus was harvested and J-Lat cells were transduced with the TxLatent library at a multiplicity of infection of 0.2. After 13 days of puromycin selection, cells were treated with DMSO (ThermoFisher Scientific, D139-1), vorinostat (500nM, SelleckChem, S1047), AZD5582 (10nM, MedChemExpress, HY-12600), prostratin (75nM SigmaAldrich, P0077), or iBET151 (75nM, SelleckChem, S2780).

### CRISPR/Cas9-mediated knockout of host genes in primary CD4 T cells

Pre-designed CRISPR RNAs (crRNAs) against ETS1 (CAAGACGGAAAAAGTCGATC, CAGAAACCCATGTTCGGGAC, and CGAGAAAGCAGTCTTTACCC), and other human gene targets, CDK6, TNFAIP3, ZNF740, SAMD12, FOXE3, ZIC5, SMC3 (Table S5) and/or HIV-1 Tat and scrambled guides (non-target control, NT) were purchased from Integrated DNA Technologies (IDT). Custom crRNAs for MALAT1 (ACTTCTCAACCGTCCCTGCA, CTGGTTCTAACCGGCTCTAG, and CCTGACGCAGCCCCACCGGTT), reported previously (76, 77), were purchased from IDT. crRNAs with superior off-target scores were selected and used for the experiment. Targeting sequences of crRNAs are provided in **Table S5**. Annealing of crRNA/tracrRNA, and preparation of CRISPR/Cas9 ribonucleoprotein complexes (RNPs), and electroporation was performed as described previously (6). Briefly, two or three crRNAs targeting different regions of the target were multiplexed for more efficient target knockout. For each electroporation, 2-3×10^6^ HIV infected primary CD4 cells were washed twice with phosphate buffered saline (PBS) at a speed of 90g for 10min then resuspended in nucleofection buffer P3 (Lonza). The resuspended cells were combined with crRNP complexes and were immediately transferred into the cuvette of the P3 Primary Cell Nucleofector Kit (Lonza; V4SP-3096) and electroporated using code CM-137 on the Lonza 4D-Nucleofector. P2 buffer was used for cells isolated from PWH and program, EH100 was used. Electroporated cells were resuspended with 200μL of prewarmed supplemented RPMI media with IL-2 (20U/mL) and IL-7 (4ng/mL) and expanded in a 37°C incubator for subsequent experiments. The cells were maintained at ∼1×10^6^ cells/mL with fresh media supplemented with IL-2 and IL-7 every 2-3 days.

### Quantification of gene expression using quantitative-PCR

RT-qPCR for HIV Gag RNA and MALAT1 RNA was performed as previously described (72). Briefly, RNA was extracted from cells harvested at three days or 2 weeks post nucleofection using the RNeasy plus kit (Qiagen) as per the manufacturer’s instructions and quantified by nanodrop. 1μg of RNA for cells from PWH and 100ng of RNA from *in vitro* primary CD4 T cells were reverse transcribed and amplified using Fastvirus (Thermo, Waltham, MA) and primer sets for HIV Gag RNA (GAG-F: ATCAAGCAGCCATGCAAATGTT, GAG-R: CTGAAGGGTACTAGTAGTTCCTGCTATGTC, GAG Probe: FAM/ZEN-ACCATCAATGAGGAAGCTGCAGAATGGGA-IBFQ). A Gene expression assay primer/probe kit from Thermo-Fischer was used for the quantification of MALAT1 (Hs00273907_s1, Thermo Fischer Scientific). Reactions were performed in 96-well plates using the Quant Studio 3 Real-Time PCR system (Applied Biosystems, Foster City, CA) real time thermocycler with a cycling parameters of 5 min reverse transcription step at 50°C, 95°C for 20 sec for Taq activation, followed by 40 cycles of 95°C (3 sec.), and 60°C (30 sec.). All qPCR reactions were performed in triplicate. Normalized relative expression levels were calculated using the Prism software version 10.1.1 (GraphPad).

### RNAseq analysis

Total RNA was harvested from nucleofected CD4 cells from PWH using a RNEasy plus kit (Qiagen). The quality of the RNA was assessed using an Agilent 2100 bioanalyzer with the RNA 6000 Pico Chip (Agilent Technologies, Amstelveen, The Netherlands). The quantity of RNA was calculated using Qubit (Thermo Fisher) and the integrity of the RNA was assessed as RNA integrity number (RIN) scores. RNAseq libraries were prepared using the KAPA Total RNA library prep kit, followed by 50bp paired end sequencing on a Nextseq2000 P2 flow cell. 70-90 million reads were obtained per sample. To analyze the data, the raw reads were first filtered by FastQC and CutAdapt to remove low quality reads. STAR version 2.7.1a was then used to align the trimmed reads to integrated human genome (GRCh38) and HIV genome (HXB2) followed by featurecounts to quantify transcripts (78). Differentially expressed genes were then identified by DESeq2 (79). Database for Annotation Visualization and Integrated Discovery (DAVID) was used for functional annotation bioinformatics microarray analysis to determine the functional enrichment (80). Bam files generated from RNASTAR were used as input for integrative genome viewer (IGV) to visualize gene transcript abundance (81).

### Immunoblotting

To extract cellular protein for western blotting, cells were lysed using RIPA buffer. Lysates were resolved on a NovexTM WedgeWellTM 4–20% Tris-Glycine gel (Invitrogen; XP04202BOX) and transferred to nitrocellulose membranes (Invitrogen; IB23002). Blocking was performed for 1h at room temperature using 5% milk in Tris-buffered saline (TBS). Immunoblotting was performed using the primary antibodies, ETS1 (Cell signaling Technology; 14069) at 1:1000, and Beta actin (abcam; ab49900) at 1:10000. Membranes were washed with 0.1% TBS-tween (TBST) 3 times for 10 min each. The following secondary antibodies were used at a 1:10000 dilution: donkey anti-rabbit IgG-HRP (Novex; A16035), and donkey anti-mouse IgG-HRP (Novex; A16011). Membranes were developed with ECLTM Prime Western Blotting detection reagents (Cytiva; RPN2232) and visualized on a BioRad Chemidoc MP Imaging System.

## Data and statistical analysis

Flow cytometry data were analyzed using FlowJo version X10.0.7r2 (FlowJo LLC; Ashland, OR, USA). Uninfected primary CD4 cells were used as a control for GFP gating for CRISPR knockout experiments. GraphPad Prism v. 10.1.1 (GraphPad; San Diego, CA, USA) was used for plotting the data. For J-Lat 10.6 cells, background GFP from unstimulated cells was subtracted from all samples prior to normalizing. All data are presented with error bars as mean ± standard deviations from at least 3 independent experiments. Statistically significant differences in provirus expression were determined by two-way ANOVA Tukey’s multiple comparisons test, with a p value <0.05 considered significant.

## Author Contributions

Conceptualization: Manickam Ashokkumar, Edward P Browne.

Data curation: Manickam Ashokkumar, Terry Hafer.

Formal analysis: Manickam Ashokkumar, Terry Hafer.

Funding acquisition: Edward P Browne, Michael Emerman.

Investigation: Manickam Ashokkumar, Edward P Browne, Michael Emerman.

Methodology: Manickam Ashokkumar, Terry Hafer, Abby Felton, Edward P. Browne.

Project administration: Edward P Browne.

Resources: Michael Emerman, David M Margolis, Nancie Archin, Edward P. Browne.

Supervision: Michael Emerman, Nancie Archin, Edward P Browne.

Validation: Manickam Ashokkumar, Edward P Browne.

Visualization: Manickam Ashokkumar

Writing – original draft: Manickam Ashokkumar, Terry Hafer, Micael Emerman, Edward P Browne.

Writing – review & editing: Manickam Ashokkumar, Terry Hafer, Abby Felton, Nancie Archin, David M Margolis, Michael Emerman, Edward P Browne.

## Acknowledgement

We thank the UNC HIV cure center, Center for AIDS Research, the UNC Flow Cytometry Core Facility, and the Fred Hutch Bioinformatics and Genomics Cores.

## Figure Legends

**Supplementary Figure S1.**
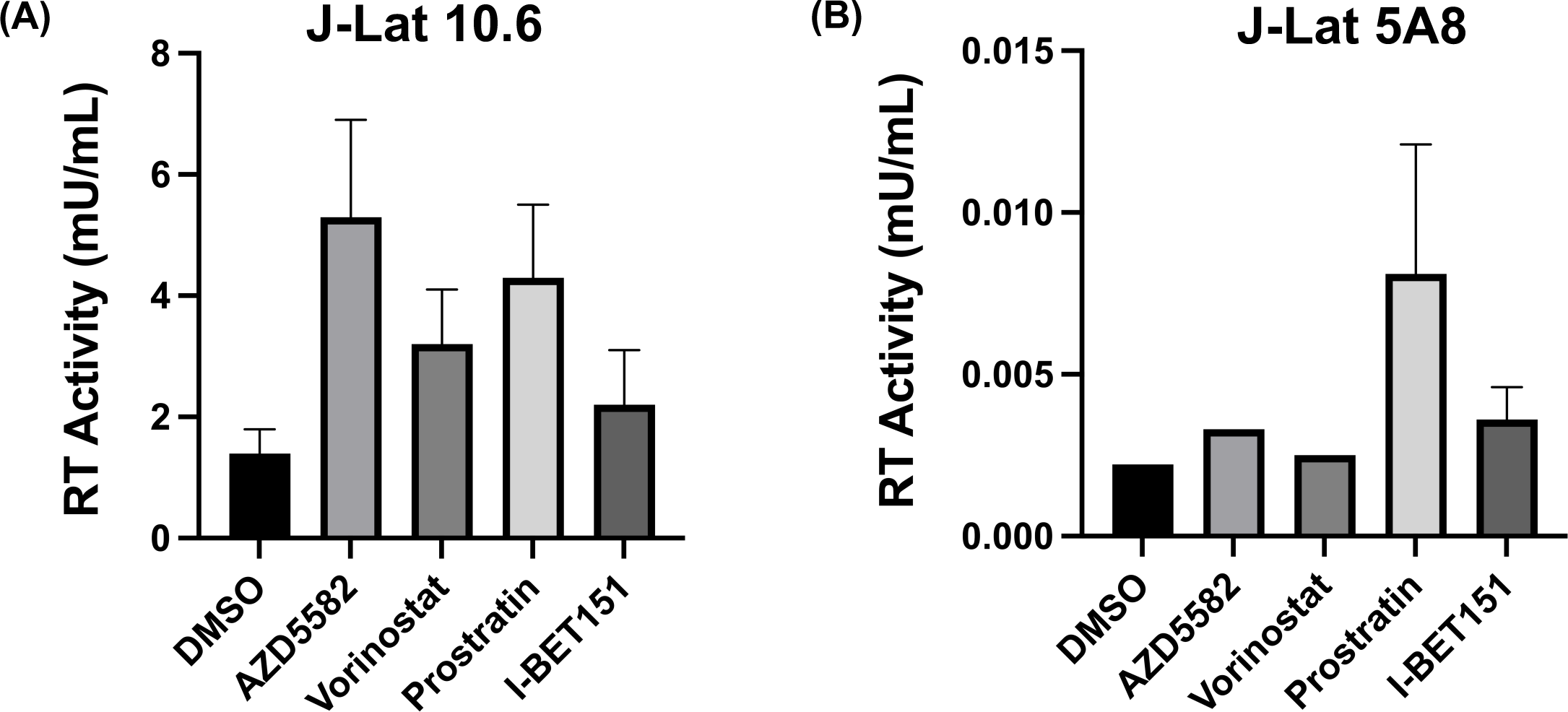
HIV-1 reverse transcriptase (RT) activity of cell free virus. Viral supernatant collected from cells at 21 days post lentiviral CRISPR transduction and stimulated for 24h with different latency reversing agents (LRAs) or DMSO control in J-Lat 10.6 (**A**) and J-Lat 5A8 (**B**) cell model was quantified for RT activity of HIV-1 proviruses expressed. Combined lentiviral transduction and LRAs treatment results in significant increase in viral reactivation.

**Supplementary Figure S2:**
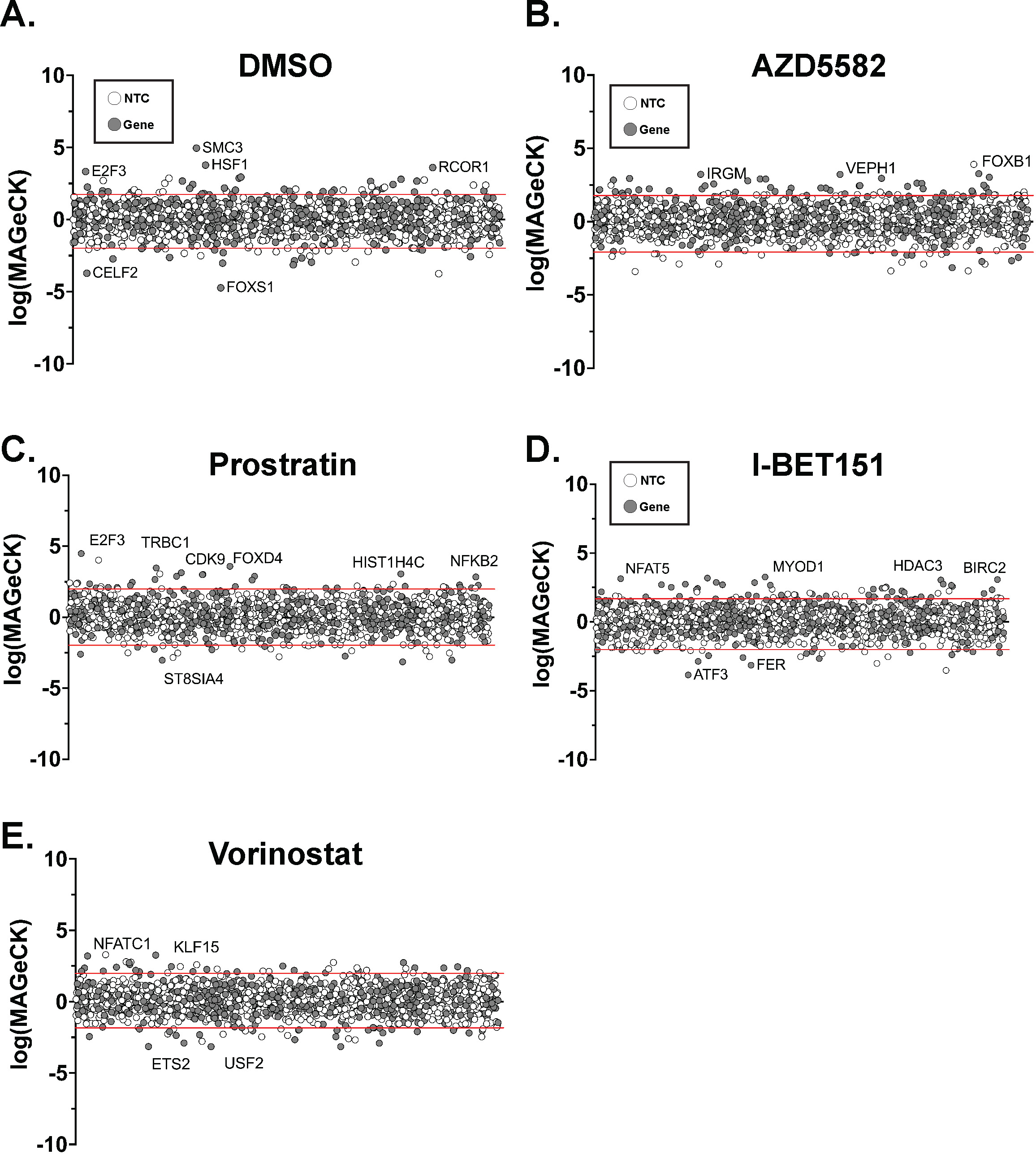
The Latency HIV-CRISPR Screen is displayed for J-Lat 5A8 cells transduced with TxLatent library and subsequently treated with DMSO (A), AZD5582 (10nM) (B), prostratin (75nM) (C), iBET151 (75nM) (D), or vorinostat (500nM) (E). Genes in the TxLatent library are randomly displayed on the x-axis, in the same order across each treatment. Synthetic non-targeting controls (SynNTCs - random groups of 8 NTCs) are shown with white circles and genes targeted by gRNAs in the TxLatent library are shown with gray circles. The -Log(MAGeCK Score) is plotted for the enriched genes and the Log(MAGeCK score) is plotted for depleted genes, and combined together to display on a single graph for each treatment. The red lines on each graph represent the two standard deviations of mean of the synNTCs: DMSO, 1.74 and -1.99; AZD5582, 1.79 and -2.09; iBET151, 1.68 and -2.01; prostratin, 1.99 and -1.97; and vorinostat, 1.98 and -1.83.

**Supplementary Figure S3.**
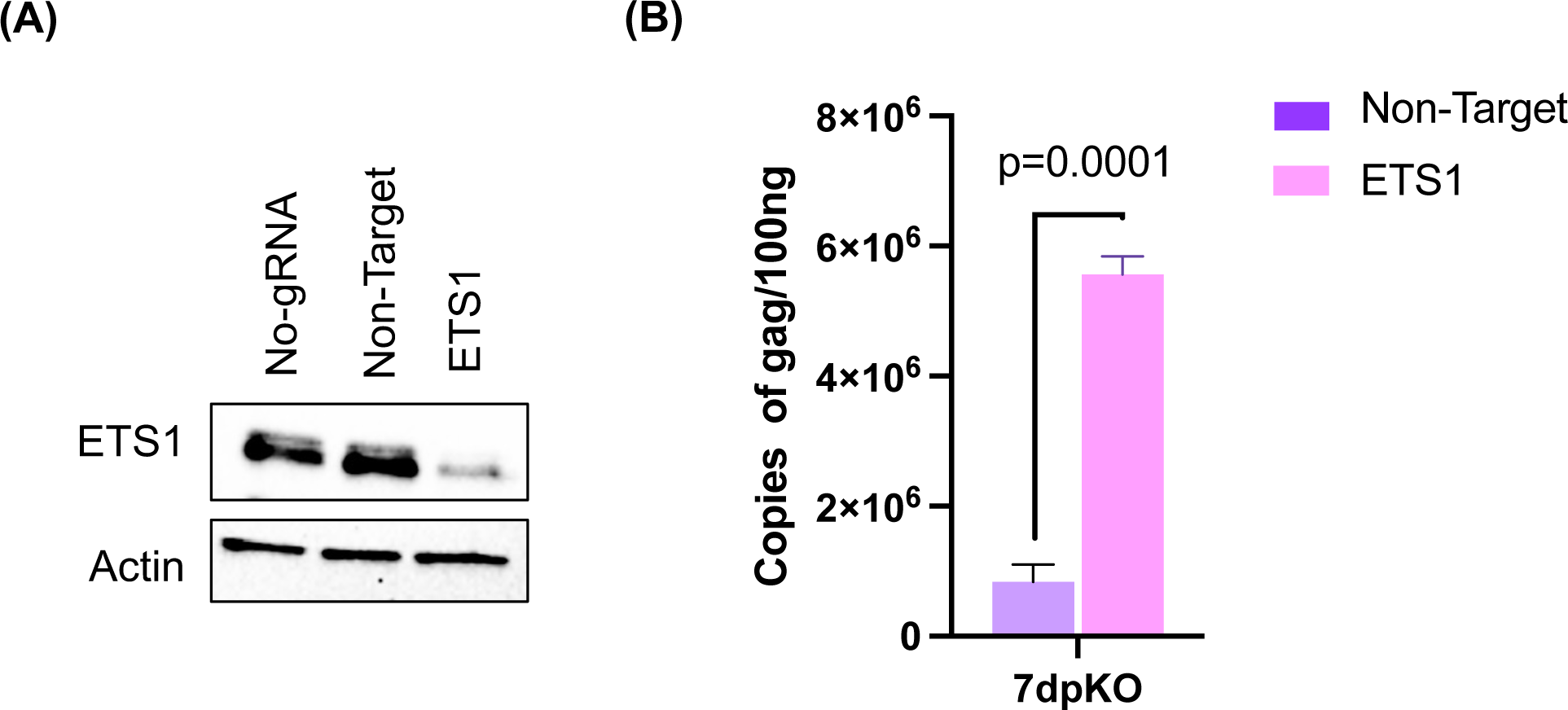
ETS1 knockout in JLat10.6 cell line model reactivates HIV. **(A).** Western blot of the ETS1-targeting or a control non-targeting (NT) gRNA nucleofected Jurkat latent cells (JLat10.6) for beta actin and ETS1 expression at one-week post nucleofection. **(B)** Relative abundance of Gag viral RNA expression in latently infected Jurkat cells nucleofected with RNPs including ETS1-targeting or NT control gRNA/Cas9 complexes. Gag viral RNA expression was normalized to input RNA. Error bars represent standard deviation, and P values displayed were determined by two-way ANOVA Tukey’s multiple comparisons test.

**Supplementary Figure S4.**
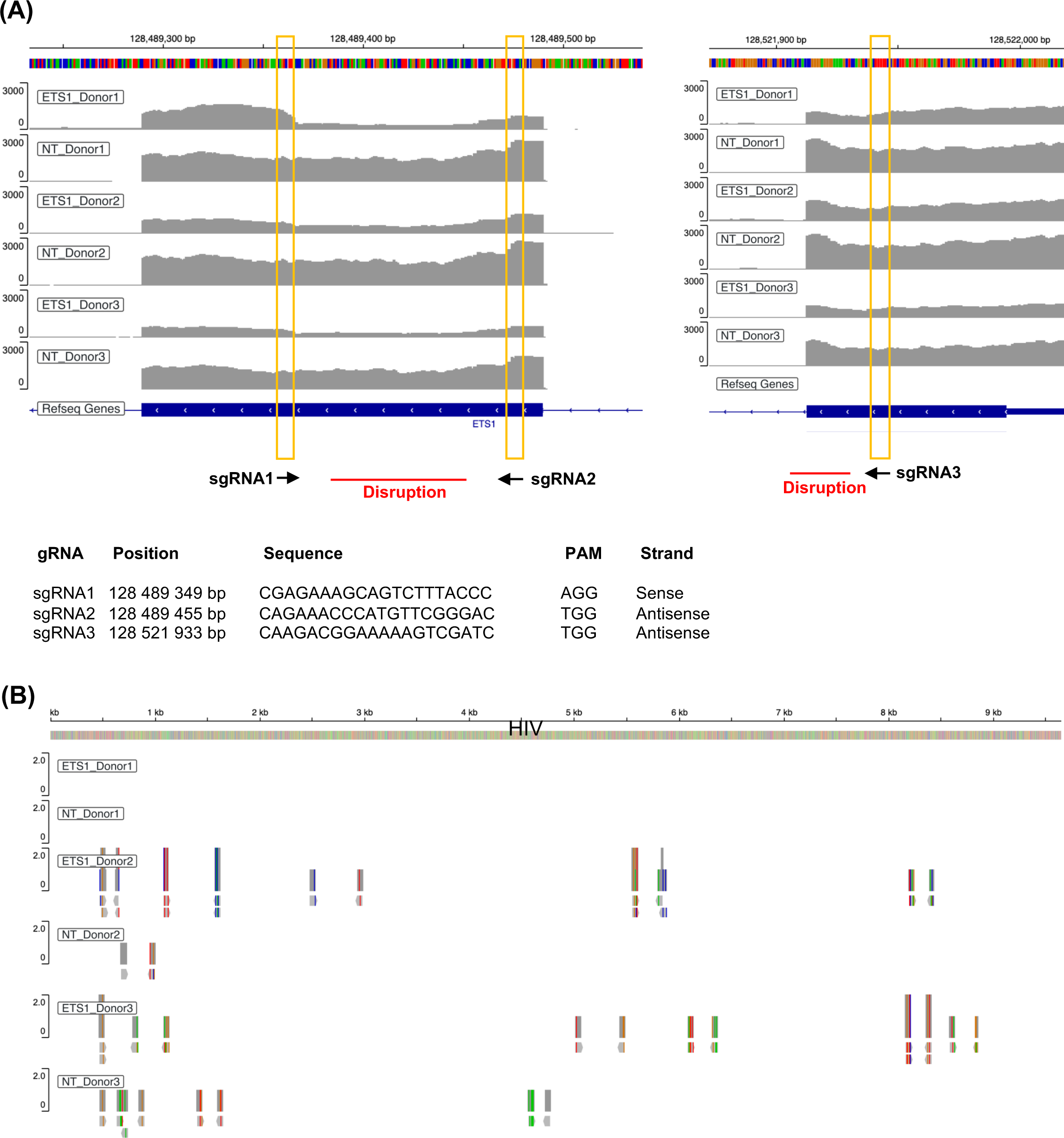
IGV display of ETS1 read disruption and HIV reads in ETS1 nucleofected CD4T cells from PWH. **(A)** ETS1 mapping reads for each of the two conditions (ETS1 and NT control) are visualized using the Integrative Genomics Viewer (IGV). The sudden change in read depth between sgRNA targeting sites is the characteristic of indels that cause larger disruption of ETS1 transcript due to CRISPR nucleofection. The read disruption is present in all the three of the ETS1 nucleofected cells, approximately half the reads and none of the reads in the NT control. Position, strand and sequence of the gRNAs for the ETS1 target is provided at the bottom. (**B**) Spliced HIV reads in ETS1 and NT control. HIV reads were undetected in sample (Donor 1).

**Supplementary table 1.** List of top hits identified in combined single multi-omic profiling of a 2D10 cell model of HIV latency.

**Supplementary table 2.** List of top hit genes screened in **Lentiviral** CRISPR Screen of TxLatent library obtained from integrated single cell multi-omics of RNA-seq and ATAC-seq of reactivated 2D10 cells with vorinostat, prostratin, AZD5582, and iBET151.

**Supplementary table 3.** Differentially expressed genes (up- and down-regulated) that correlate with HIV RNA levels in ETS1 depleted CD4 T cells isolated from ART suppressed PWH.

**Supplementary table 4.** List of genes and corresponding guides that comprise the TxLatent Library.

**Supplementary Table 5:** List of gRNAs targeting different transcripts/transcription factors in primary latent cell model.

